# A Minimally Invasive Technique for Serial Intraosseous Perfusion Measurements in the Murine Tibia Using Laser Doppler Flowmetry

**DOI:** 10.1101/708453

**Authors:** Nicholas J Hanne, Elizabeth D Easter, Sandra Stangeland-Molo, Jacqueline H Cole

## Abstract

In biomedical and preclinical research, the current standard method for measuring blood perfusion inside murine bone, radiolabeled microspheres, is a terminal procedure that cannot be used to monitor longitudinal perfusion changes. Laser Doppler flowmetry (LDF) can quantify perfusion within the proximal tibial metaphysis of mice *in vivo* but requires a surgical procedure to place the measurement probe directly onto the bone surface. Sustained inflammation for over a month following this technique was previously reported, and previous studies have used LDF as an endpoint-only procedure. We developed a modified, minimally invasive LDF procedure to measure intraosseous perfusion in the murine tibia without stimulating local or systemic inflammation or inducing gait abnormalities. This modified technique can be used to measure perfusion weekly for up to at least a month.

- Unlike previous endpoint-only techniques, this modified LDF procedure can be performed weekly to monitor serial changes to intraosseous perfusion in the murine tibia
- The modified LDF technique utilizes a smaller, more localized incision to minimize invasiveness and speed recovery

## Method Details

### Background

The current standard method for assessing functional blood flow within bone in rodents uses microspheres labeled with fluorescent or radioactive tags that are introduced to the vascular supply, and then the amount of microspheres in tissue samples taken from the animal *ex vivo* are quantified [1,2]. However, this procedure is a terminal one and cannot be used to assess serial changes to intraosseous blood flow. Laser Doppler flowmetry (LDF) has been validated as an accurate technique to quantify *in vivo* perfusion within the murine proximal tibial metaphysis, but signs of local inflammation for over a month after the procedure were reported [3]. Inflammation stimulates angiogenesis, which could alter subsequent perfusion measurements [4]. Similarly, inflammation may affect nearby bone remodeling via cellular signaling [5] or alter limb loading via gait abnormalities [6], both of which could confound study designs focused on bone tissue outcomes. We have modified the LDF technique to minimize invasiveness so that it can be used for serial measurements of bone perfusion at weekly intervals in the same mice without causing inflammation [7].

### Modified LDF Procedure

This protocol was approved by the Institutional Animal Care and Use Committee at North Carolina State University. In this study, the LDF procedure was performed on eighteen 14-week-old male C57Bl/6J mice (The Jackson Laboratory, Bar Harbor, ME). Mice should be fasted 6-8 hours prior to the procedure. The LDF probe is sensitive to movement, temperature, and light, so care should be taken to ensure consistent conditions across procedures. The LDF monitor should be regularly calibrated with a probe flux standard. Follow the manufacturer’s directions to calibrate the probe. Calibrate between studies, not between LDF procedures.

#### LDF Monitor and Software Preparation

1. Connect the LDF monitor to a computer. Turn on the LDF monitor and open the data acquisition software provided by the manufacturer. In the signal processing settings, reduce the bandwidth to one appropriate for the slower blood speeds in rodent bone, about 0.4-1 mm/s [3], by selecting a lower cutoff frequency for the low-pass filter (e.g., 3 kHz) [8], as suggested by the manufacturer. *Note: Perfusion data captured with different bandwidth settings should not be compared*.
2. Clean the LDF probe with 70% ethanol using a sterile cotton swab and allow to air dry.

#### LDF Measurement Procedure

1. Anesthetize mouse in an induction chamber with isoflurane (3-5%) in pure oxygen. *Note: Different gas mixtures (i.e., compressed air) can be used instead of pure oxygen but may alter intraosseous blood supply. Perfusion data should not be compared for different anesthesia conditions*.
2. Shave the fur over the anterior and medial surfaces of the proximal tibia and knee joint.
3. Place the mouse supine on a heated surgical pad, and tape down the hind paws so that the medial surface of the hindlimb is accessible. Maintain anesthesia with isoflurane (1.5-2%). Maintain rectal temperature at 37°C with a feedback-controlled heating system. Apply ophthalmic ointment to the eyes to prevent corneal drying during the procedure.
4. Disinfect the incision site three times with povidone-iodine followed by 70% ethanol.
5. Gently palpate the anterior surface of the proximal tibia with forceps to find the most proximal surface over the anteromedial surface of the bone that does not have soft tissue beneath the skin. Position the limb (re-tape limb if needed) so that this site is easily accessible (Figure 1A).
6. Using a #11 or other preferred small scalpel blade, make a 2-5-mm long sagittal skin incision directly over the proximal tibia, starting approximately 3-4 mm below the knee and moving distally (Figure 1B). Use a sterile cotton-tipped applicator to absorb fluid at the incision site. *Note: This incision should not cause excessive bleeding. If blood pools in the incision site, making probe placement difficult, firmly hold a sterile cotton-tipped applicator or gauze for 15-30 seconds until bleeding stops*.
7. Using forceps, gently retract any soft tissue covering the tibia and palpate the tibia to ensure no soft tissue is covering the bone. Using the edge of the scalpel blade, gently scrape a small window in the periosteum that is approximately the same size as the LDF probe tip (0.8 mm diameter). Probe the bone surface gently with the scalpel to confirm the periosteum was removed. Before removal the surface will feel more compliant when compressed, and after it is removed it will feel harder.
8. Using a micromanipulator, position the LDF probe tip in the periosteum-free window, placing it firmly against and perpendicular to the exposed bone surface (Figure 2). Pushing the probe with excessive force into the bone can lower tissue perfusion measurements, so position it just firmly enough to prevent bone movement or probe slipping during recording. *Note: The LDF probe is sensitive to motion. Do not touch the LDF probe or fiber optic cable while recording perfusion*.
9. Record perfusion with the LDF monitor’s software. We capture 30-second perfusion readings, which are sufficient for a stable average measurement, but longer or shorter readings could be recorded, if desired. *Note: Perfusion readings under 5 PU indicate the probe is not placed perpendicular to the medullary cavity, and readings over 30 perfusion units (PU) likely indicate the probe slipped off the bone and is recording in soft tissue. A noisy signal (greater than 5 PU variation) suggests that either the probe is not pressed firmly against the bone, the probe is moving, or the probe is not perpendicular to a flat bone surface. In any of these cases, reposition and secure the probe until resolved. Large spikes in the signal likely indicate limb movement; sufficient anesthesia depth should be confirmed, and the hindlimb can be taped down above the knee if additional security is needed*.
10. After obtaining a good reading, remove and reposition the LDF probe, and record again. *Note: We recommend repositioning the probe at least one time to ensure consistent placement over the bone. If the two readings are dissimilar (greater than 5 PU difference), then more readings should be taken and averaged*.
11. Remove the probe. Pinch the incision closed with forceps, and apply a small amount of tissue adhesive to the incision. Continue to hold the incision closed for 15 seconds, being careful to avoid contacting the glue with the forceps. *Note: Tissue adhesive should only be applied to the outside of the incision, not within the incision. The glue dries rapidly, and only a small amount is needed to close a small incision*.
12. Apply 2% lidocaine cream and triple antibiotic ointment to the incision site.
13. Remove the tape from the hindlimbs, remove the rectal probe, and turn off the isoflurane. Move the mouse to a heated recovery cage until ambulatory. Return to normal cage and monitor for signs of discomfort twice a day for at least 48 hours and longer if necessary.

**Figure 1:**
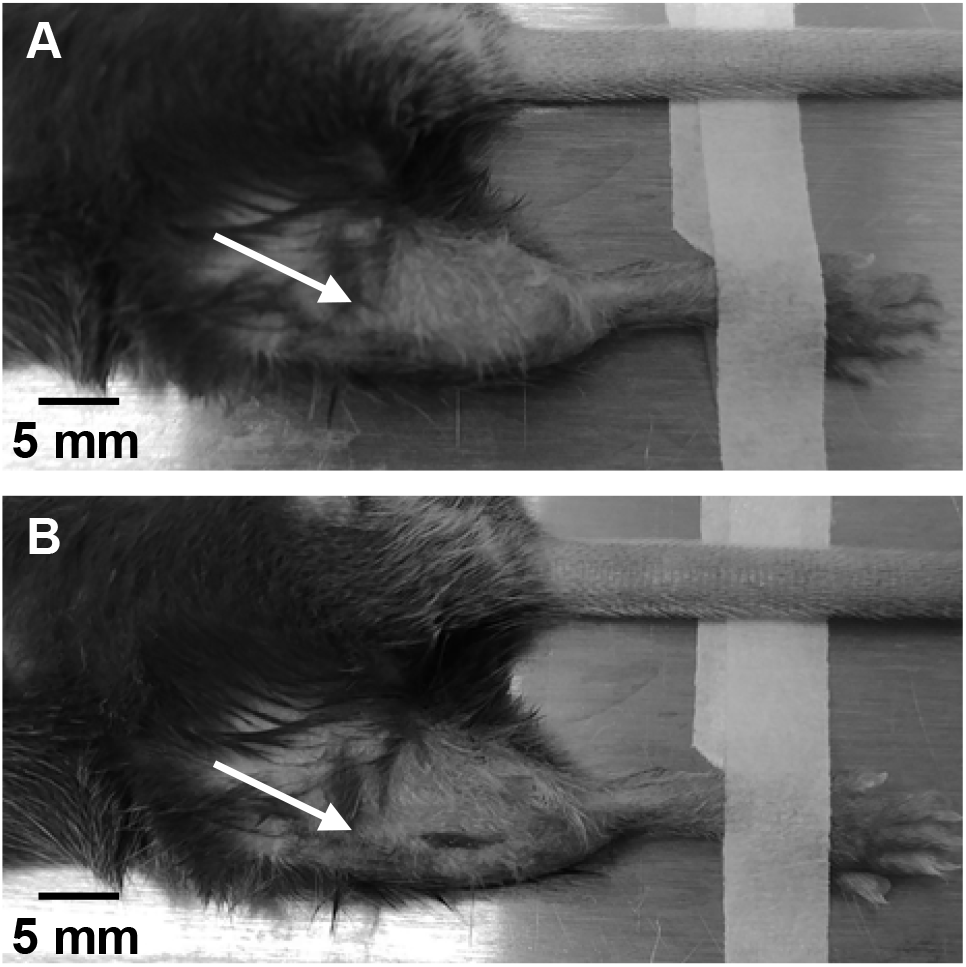
Image showing the measurement site on the right hindlimb A) before and B) after the incision is made. The arrows point to the approximate position of the right knee.

**Figure 2:**
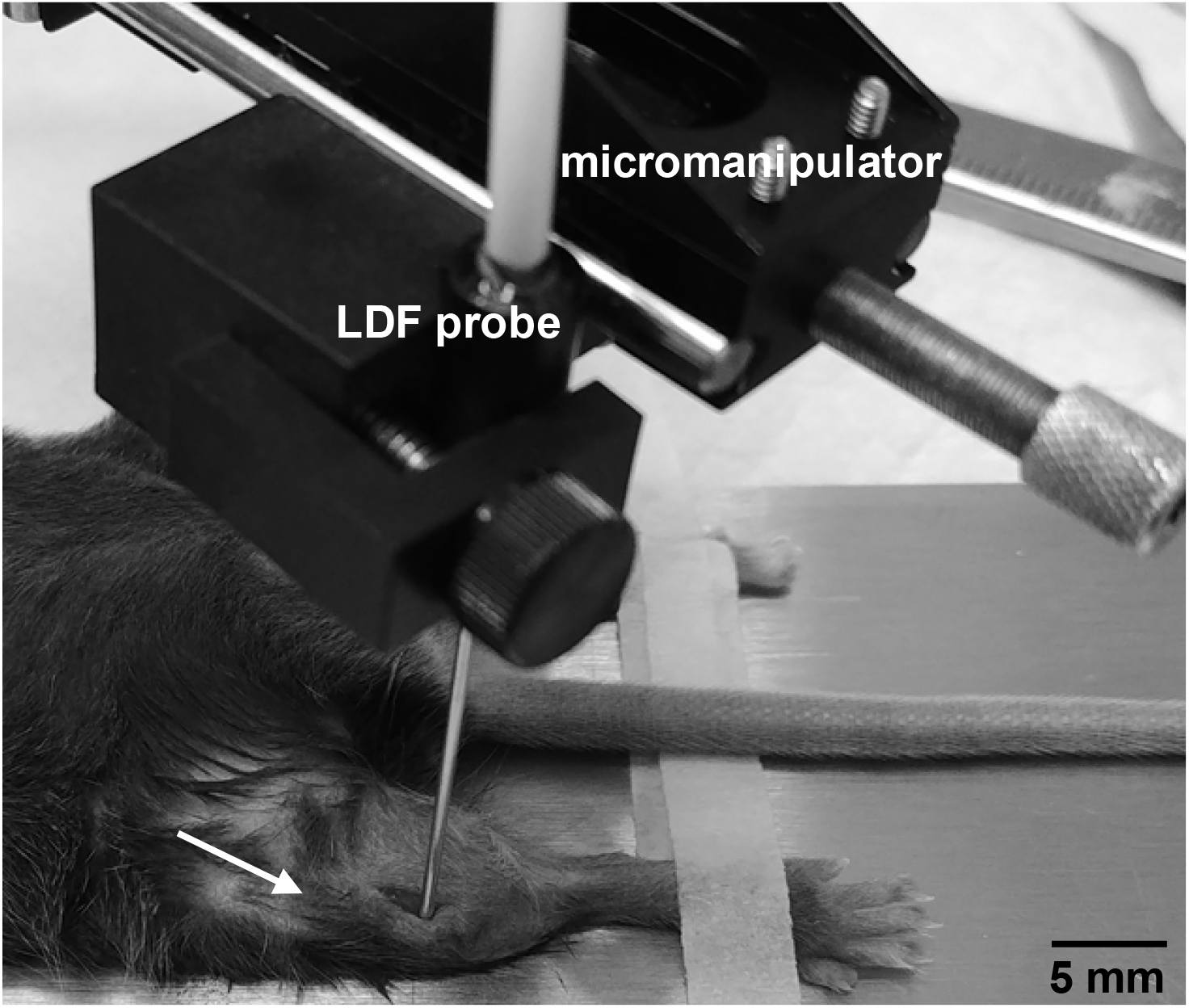
Image showing probe placement during the modified laser Doppler flowmetry technique on the right hindlimb. The arrow points to the approximate position of the right knee.

#### Data Processing

1. Select individual readings in the LDF monitor software, avoiding spikes if possible. Export the mean perfusion and length of the reading, which can be either in time units or number of points.
2. Tibial perfusion for each mouse is calculated as the weighted mean of each of the readings for each mouse:

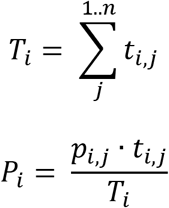

where *T_i_* is the total length of all *n* readings for each mouse (*i*), *t_i,j_* is the length of each reading (*j*) for that mouse, *P_i_* is the weighted mean perfusion over all readings for that mouse, and *p_i,j_* is the mean for each perfusion reading for that mouse.

### Validation

Our modified LDF technique is based on a procedure that has been proven to accurately measure perfusion in the murine tibia, demonstrated by comparing LDF readings to controlled blood flow rates through *ex vivo* bone samples using a syringe pump [3]. Greater cortical thickness will have a lower LDF perfusion measurement, but small variations in cortical thickness do not greatly affect perfusion readings [3]. Therefore, LDF readings should only be compared when cortical thickness is not expected to be significantly different, whether with age, sex, mouse strain, treatment, or other conditions.

We validated that this modified technique can be used serially in the same mice without inducing inflammation or gait abnormalities [7]. We performed the modified technique in two groups of male C57Bl/6J mice starting at 12 weeks of age. One group received weekly tibial perfusion measurements for four weeks (n=10, *Repeated*), while the other group only had tibial perfusion measured at the fourth week (n=8, *Endpoint*). We found no difference in tibial perfusion between the Repeated and Endpoint groups at the end of the study (t-test p = 0.92, Figure 3), demonstrating that repeated LDF procedures do not affect subsequent measurements, at least up to four weeks. We also showed that the modified technique does not alter hindlimb gait patterns, induce localized inflammation at the incision site, or cause detectible increases in serum concentrations of inflammatory marker interleukin-6 when performed weekly in the same mice for four weeks.

**Figure 3:**
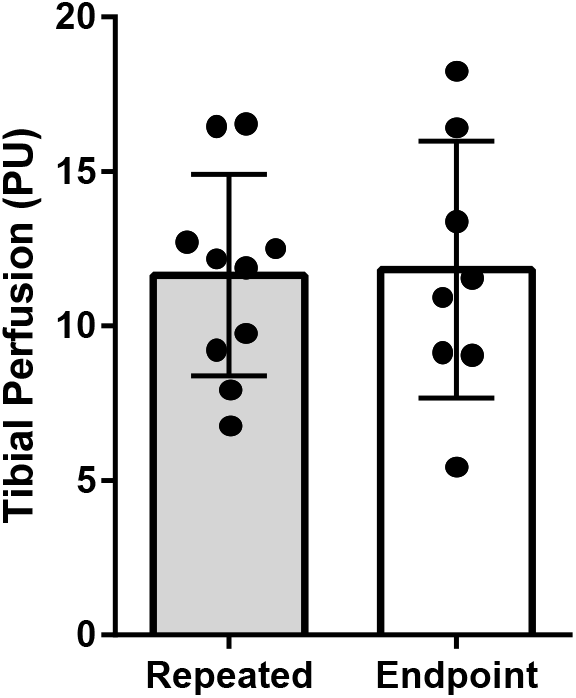
Perfusion measured in the proximal tibia was similar between mice that received a weekly laser Doppler flowmetry procedure for four weeks (*Repeated*) and mice that received the procedure only once at the final timepoint (*Endpoint*).

While our modified LDF procedure has only been validated for serial bone perfusion measurements in young adult male C57Bl/6J mice just after skeletal maturity, it may also be useful in other rodent populations. We have applied this technique to serial measurements over four weeks in 24-week-old male and female C57Bl/6J mice and to endpoint measurements in aged male Brown Norway rats at 34 months of age. Other studies have performed endpoint-only LDF procedures in older mice (17 months of age) [3], female mice [3], and rabbits [9]. A full validation should be performed before performing this serial measurement procedure in different populations to ensure the animals do not respond differently to weekly procedures. Because age, sex, and mouse strain affects LDF perfusion measurements [3], studies involving this technique should be designed with caution to avoid comparisons across different populations.

